# A widespread SCC*mec*-located gene cluster protects methicillin-resistant *Staphylococcus aureus* against toxic polysulfides

**DOI:** 10.64898/2026.01.28.702267

**Authors:** Verena Wiemann, Jan-Samuel Puls, Tomohisa Sebastian Tanabe, Jan-Martin Daniel, Sharmila Sekar, Simon Heilbronner, Tanja Schneider, Christiane Dahl, Fabian Grein, Thomas Fließwasser

## Abstract

The genus *Staphylococcus* contains important human commensals and pathogens, including methicillin-resistant *Staphylococcus aureus* (MRSA), which is a frequent colonizer of humans and a leading cause of healthcare-associated and life-threatening infections. While its virulence and pathogenicity have been extensively studied, factors driving the colonization and distribution of MRSA as a pathobiont are less understood. Here, we report on a *cst* sulfide detoxification gene cluster located on SCC*mec,* the antibiotic resistance-mediating genetic element of MRSA. Bioinformatic analyses revealed a heterogeneous distribution of *cst* clusters in staphylococcal genomes and that many clinically relevant SCC*mec* types introduce an additional *cst* cluster (*cst2*) to MRSA. While the canonical *cst* cluster (*cst1*) consists of the five genes *tauE*, *cstR*, *cstA*, *cstB*, and *sqr*, most staphylococcal *cst* clusters, including the SCC*mec*-located *cst2*, lack the *sqr* gene, which encodes for a sulfide:quinone reductase responsible for the initial step of sulfide detoxification. Growth experiments with a diverse set of representative *Staphylococcus* strains, *cst*-deletion mutants, and complementation with *cst*-containing plasmids demonstrated that the *cst* cluster enables *sqr*-independent polysulfide-detoxification. Furthermore, the additional *cst2* cluster confers high polysulfide tolerance to MRSA, providing the pathogen with a unique advantage in polysulfide-rich environments. Using serial passaging co-cultivation experiments with methicillin-sensitive *S. aureus* (MSSA) strains, we demonstrated that in the presence of polysulfides *cst2*-containing MRSA can invade an established MSSA population and outperform the occupying resident in direct competition. Overall, our findings indicate that polysulfides are critical stress factors for staphylococci, potentially contributing to the spread of *cst2*-containing SCC*mec* and MRSA.

**Importance:** Methicillin-resistant Staphylococcus aureus (MRSA) is one of the most prevalent human pathogens responsible for millions of life-threatening infections worldwide. It acquires antibiotic resistance through the genetic element SCC*mec*, which contains the characteristic *mecA* gene that renders the organism resistant to most classes of β-lactam antibiotics. Besides *mecA* and accessory gene complexes necessary for the transfer of SCC*mec* and phenotype manifestation, the genetic element also contains prominent gene clusters with unknown functions. Here, we report on a (poly-)sulfide-detoxification gene cluster (*cst2*) present on SCC*mec* that provides MRSA with a unique advantage in environments containing polysulfides – highly reactive intermediates of sulfide oxidation naturally occurring as microbial stressors on mucosal surfaces inside the human body. We demonstrate that in the presence of polysulfides, *cst2* enables MRSA to outperform non-MRSA in direct competition, thus supporting the invasion and proliferation of this pathogen independent of its antibiotic resistance.

## Introduction

Methicillin-resistant *Staphylococcus aureus* (MRSA) is one of the most prevalent human pathogens, responsible for millions of life-threatening infections worldwide (1). It acquires resistance to beta-lactam antibiotics through the staphylococcal cassette chromosome *mec* (SCC*mec*), a mobile genetic element. SCC*mec* contains the *mec* gene complex for resistance, the *ccr* gene complex responsible for genomic integration, and variable joining regions. Different SCC*mec* types (I-XV) and subtypes are defined by order, length, and sequence of these core components. SCC*mec* types I-III are common in hospital-acquired MRSA (HA-MRSA), while types IV-V are prevalent in community-acquired MRSA (CA-MRSA). Livestock-associated MRSA (LA-MRSA), particularly lineage ST398, often carries SCC*mec* types IVa or Vc (2).

Understanding MRSA’s spread is key to combating it. *S. aureus* frequently colonizes human mucosal surfaces, mainly the nose and the gut, where it competes with the host microbiota, especially with other staphylococci (3, 4). This competition is largely influenced by nutrient availability and stressors arising from the host and other bacteria, and it is still unclear how MRSA colonizes these environments (3). One relevant stressor stems from bacterial mucin degradation. Here, the metabolization of sulfated sugar moieties and L-cysteine residues eventually produces sulfide, which is well-known for its toxicity (5). Notably, this results in high sulfide concentrations of up to 0.4 mM in the nose (6) and 0.3 - 3.4 mM in the gut (7, 8). Under oxic conditions, which are present on these mucosal surfaces (9), sulfide undergoes rapid oxidation to form inorganic polysulfides (referred to as polysulfides from here on), toxic intermediates that ultimately react to form elemental sulfur (10–12). Interestingly, sulfide production and, consequently, polysulfide generation are further enhanced during periods of dysbiosis and infection, when the mucin layer is rapidly degraded (5). Thus, human-associated staphylococci, and in particular pathogenic species like *S. aureus*, have to deal with these highly toxic sulfur compounds (13).

It is known that *S. aureus* possesses a sulfide-detoxifying gene cluster *cst,* which encodes enzymes for the conversion of sulfide to less toxic sulfite (14). The *cst* cluster consists of genes for a sulfite transporter (*tauE*) (15), a sulfur transferase (*cstA*) (16), a sulfur dioxygenase (*cstB*) (17), and a sulfide:quinone oxidoreductase (*sqr*) (18), which are all regulated by the transcriptional repressor CstR (19, 20). According to the current model, SQR and CstB stepwise oxidize cytoplasmic sulfide to thiosulfate (Fig. S1). Then, CstA transfers the sulfane sulfur of thiosulfate to a cellular acceptor and releases sulfite, which is exported out of the cell by TauE (17, 16).

Although the biochemical properties of the *S. aureus cst* gene products are well-understood, it is currently unclear how other staphylococci cope with sulfide stress. Beyond that, the ecological role of the cluster in staphylococci is virtually unknown. Accordingly, the relationship between (poly-)sulfide stress, detoxification, pathogenesis, and niche colonization of staphylococci, particularly MRSA, is not yet understood. Strikingly, it was noted before that some MRSA strains appear to harbor a duplicate of *cst* in proximity to the resistance-conferring *mecA* gene (17). However, no substantial investigation into this phenomenon has been performed so far.

## Results

### Distribution of the *cst* gene cluster in staphylococci

We first determined the distribution of the *cst* gene cluster in the genus *Staphylococcus*. The limited annotation quality of sulfur-metabolism-associated genes and the structural similarity of respective proteins despite distinct functionality (21) prompted us to use HMSS2, a recently developed tool for the accurate detection of sulfur metabolism proteins (22). HMSS2 was extended to identify *cst* clusters in proximity to *mecA*, with core clusters labeled *cst1* and additional clusters labeled *cst2.* We found *cst1* to be widely distributed, but not highly conserved among staphylococci (Fig. 1). The species boundaries roughly defined the presence of the gene cluster. However, considerable intraspecies variation was observed, e.g., in *S. epidermidis* (16/30 strains with *cst1*). Notably, we found the *sqr* gene exclusively in *S. aureus* and additionally on a plasmid, which is present in a few *S. saprophyticus* strains, raising questions about its role in *cst1* functionality (17). The *cst2* cluster was also common in other species and often SCC*mec*-associated, as expected from the genus-wide mobility of the genetic element (23). Exceptions were the MSSA strain *S. aureus* ATCC 29213 and some plasmid-harboring *S. saprophyticus* and *S. pasteuri* strains (Table S1 and Fig. S2). We also found *mecA*-carrying strains without *cst2*, demonstrating that not all SCC*mec* types harbor the cluster.

**Fig 1.**
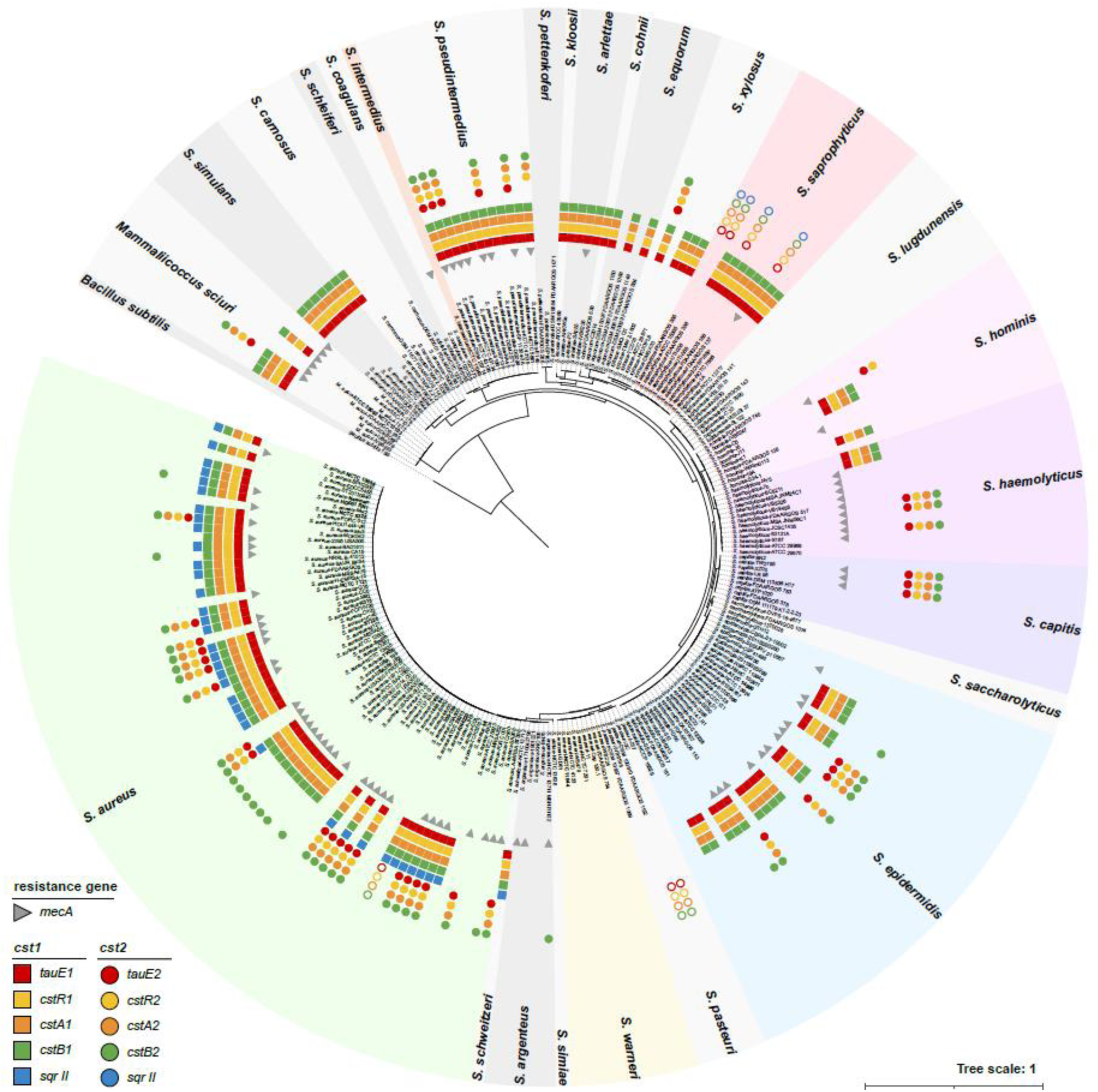
Distribution of *cst* gene clusters and *mecA* in the genus *Staphylococcus*. The *cst* genes and *mecA* gene were identified using HMSS2 (22), searching the complete genome of each strain. The genes are mapped to the maximum likelihood phylogeny of 229 strains from 26 species and *B. subtilis* 168 as outgroup. See S1 Table for a comprehensive list of all strains. The *cst2* genes that were found to be plasmid-located are represented as unfilled circles using their respective color code.

### Prevalence of the *cst2* gene cluster on SCC*mec*

To characterize the *cst2* distribution in detail, we analyzed MRSA reference strains (24–26) and epidemiologically relevant strains (27, 28), examining the spread of *cst2* across various SCC*mec* types (Table S2). We found the *cst2* gene cluster in the important HA-MRSA-associated SCC*mec* types I-III. In contrast, strains of the most prominent CA-MRSA-associated SCC*mec* type IV featured either no *cst2* cluster or a truncated version (only the gene *cst2B* in type IVa). Furthermore, the LA-MRSA-associated SCC*mec* type Vc strains all contained full-length *cst2*, while the other SCC*mec* subtypes Va and Vb had no *cst2*. Additionally, SCC*mec* types VIII, X, XIV, and XV contained a complete *cst2* cluster, whereas types VI, VII, IX, and XI-XIII lacked it. In addition, we did not find any evidence for a SCC*mec*-located *sqr* gene (Fig. S3).

We visualized the phylogenetic relationships between the genomic *cst1* clusters of staphylococci and the SCC*mec*-located *cst2* clusters of MRSA (Fig. 2A). Notably, all *cst2* clusters were distinct from *S. aureus cst1,* ruling out intraspecific duplication as the origin of the SCC*mec*-located clusters. Furthermore, we found the SCC*mec*-located *cst2* clusters separated into two groups: Group A *cst2* from SCC*mec* types II, III, VIII, XIV, and XV, and Group B *cst2* from SCC*mec* types I, V, X, and the truncated type IVa. Within each group, sequences are substantially conserved (≥ 99.91% identity in Group A, > 94.36% in Group B), but intergroup identity is approx. 72%, indicating two different origins of the clusters. While some genomic *cst1* clustered with Group B, and are thus potential origins of SCC*mec* integration, we found no genomic *cst1* close to Group A, rendering the origin of that *cst2* currently elusive.

**Fig 2.**
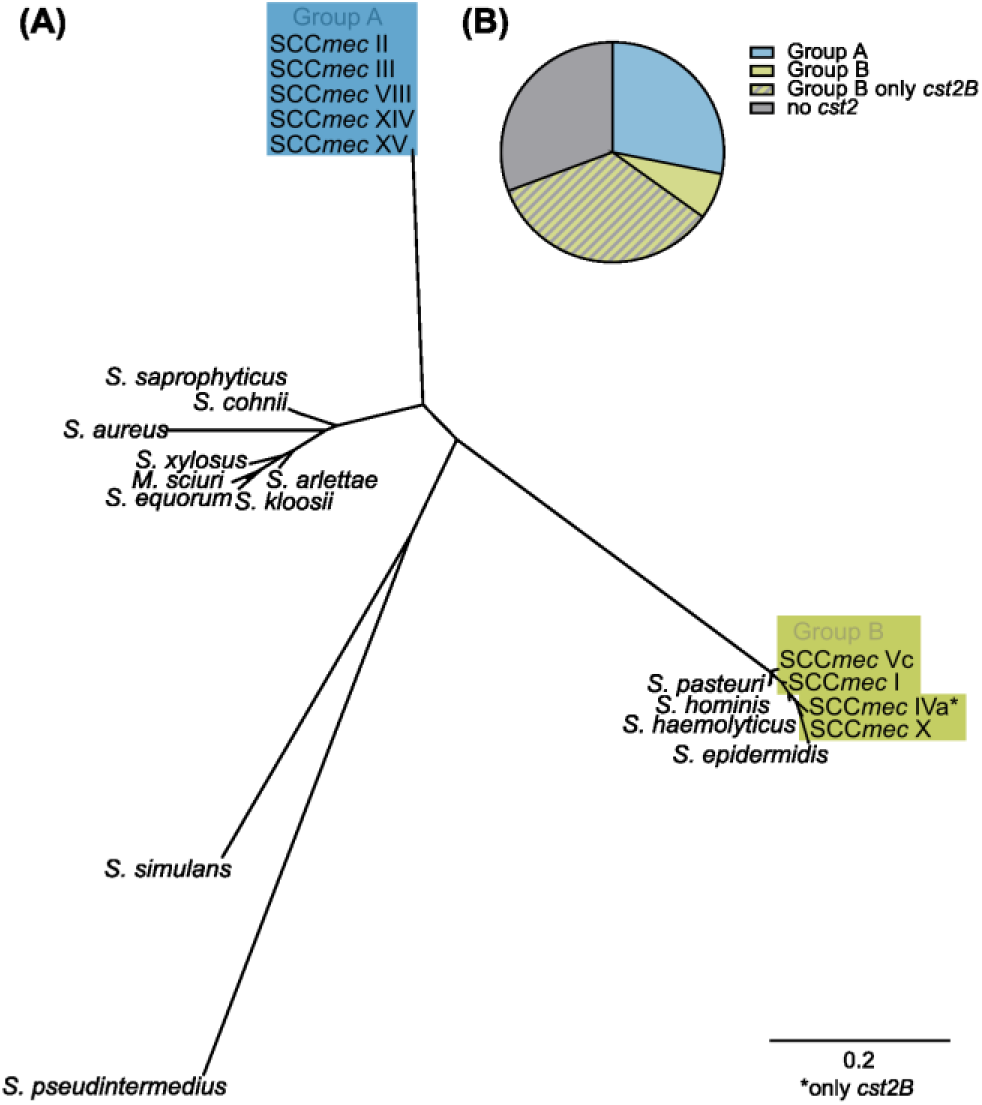
Genetic analysis of the *cst2* cluster in MRSA. (A) Radial phylogram of representative *cst1* clusters in staphylococci, and all *cst2* clusters identified in MRSA. The *cst2* groups that were identified as conserved across the diverse SCC*mec* types are indicated as Group A and Group B. (B) Relative abundance of Group A, full-length or shortened Group B *cst2* clusters (n = 282, 68, and 343, respectively) in complete MRSA genomes (n = 999) found in the NCBI *S. aureus* complete genome database.

We further investigated the prevalences of the *cst2* gene cluster Groups A and B within MRSA by screening the NCBI complete genome database. Over one-third of all complete MRSA genomes featured a full-length *cst2* gene cluster, highlighting the widespread presence of *cst2* among MRSA strains. Out of 999 complete *mecA*-containing genomes (Table S3), 282 (28.23%, Table S4) contained a Group A *cst2* and 68 (6.81%, Table S5) contained a full-length Group B *cst2* (Fig. 2A). Additionally, 343 (34.33%, Table S6) strains contained the truncated version of Group B *cst2*.

### Impact of sulfide and its oxidation intermediates on *S. aureus*

The prevalence of *cst2* in multiple SCC*mec* types suggested that MRSA benefits from the additional gene cluster. The heterogeneous distribution of the *cst1* gene cluster in staphylococci provided further evidence that it serves as a niche adaptation with relevance to survival specifically in sulfide-rich environments. Thus, we investigated the ecophysiological role of the *cst1* cluster in *Staphylococcus* species to understand how an additional *cst* cluster may enhance MRSA fitness. Previous work had shown that NaSH-containing media decreased *S. aureus* growth and that individual *cst*-encoded components partially counteracted this effect (20). To study the effects of a physiologically relevant sulfide concentration (1 mM) on the growth of common laboratory MSSA strain RN4220, we generated a Δ*cst1* mutant of the strain and compared it to the wild type. Although both strains showed a prolonged lag phase under 1 mM NaSH, the prolongation was substantially more pronounced in the deletion mutant (Fig. 3A). Notably, a considerable increase in OD600 occurred shortly after the addition of NaSH, even without bacterial cells (Fig. 3B), suggesting a spontaneous chemical reaction. Given the oxic conditions and circumneutral pH of the medium, we suspected that sulfide was being oxidized to polysulfides, which would eventually react to form highly light-refracting elemental sulfur (10, 29). Due to the reductive nature of sulfide, elemental sulfur can be reduced back to polysulfides, resulting in a continuous sulfide-polysulfide-sulfur interconversion (30). Additionally, sulfide can be oxidized to thiosulfate and sulfite under oxic conditions (Fig. 3C). This complexity of sulfide oxidation dynamics led us to identify sulfur compounds in our LB NaSH medium that may act as growth inhibitors. Therefore, we quantified polysulfides, sulfide, thiosulfate, and sulfite by HPLC, and the insoluble elemental sulfur by cyanolysis at different time points after the addition of NaSH to the medium (Fig. 3D).

**Fig 3.**
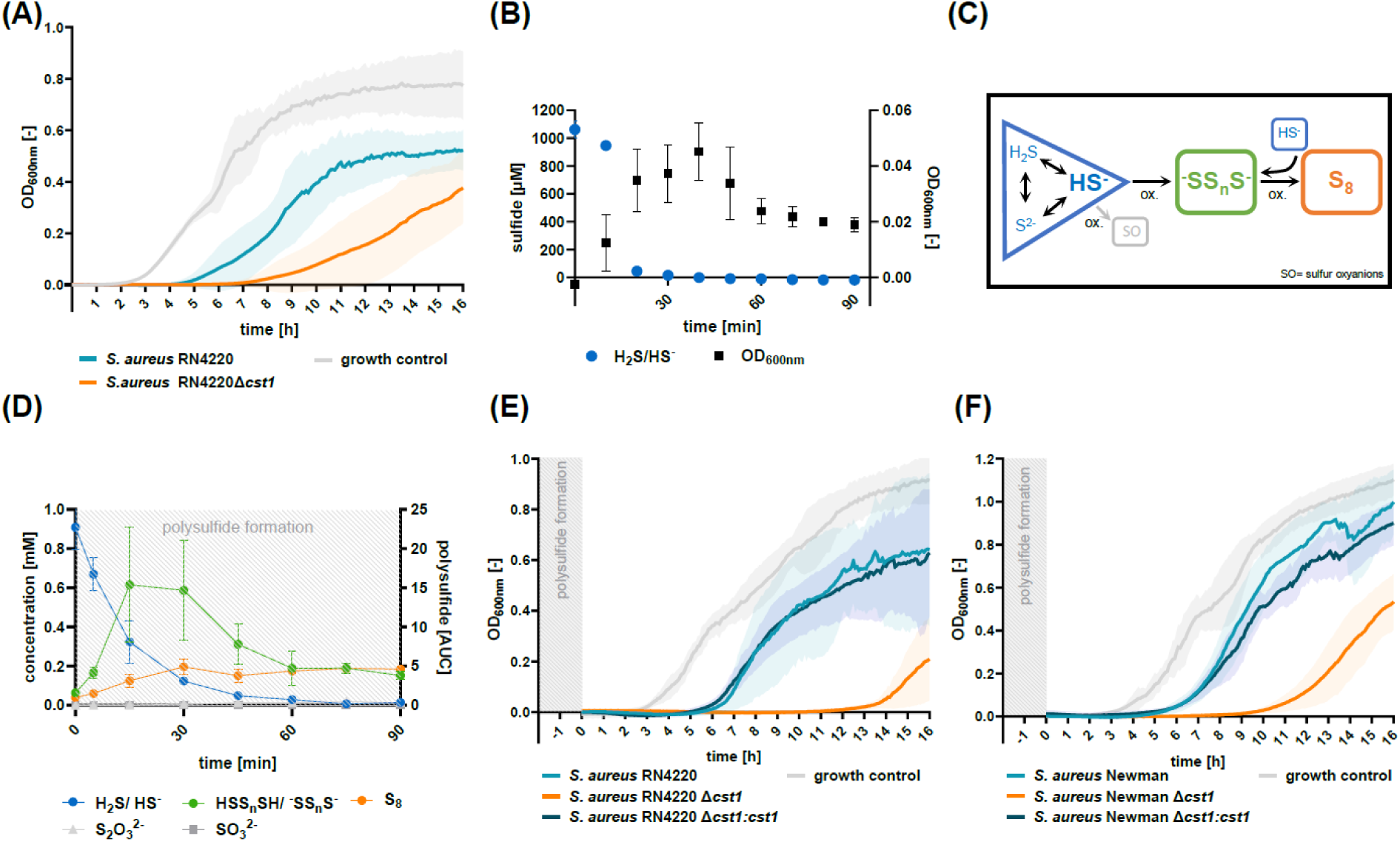
Growth behavior of *S. aureus* wild type and Δ*cst1* mutants under NaSH/polysulfide stress. (A) Growth curves of the strains RN4220 and RN4220 Δ*cst1* in LB NaSH (1 mM) medium. Plotted are the mean values of three independent, biological replicates. Light-hued areas illustrate the standard deviations (SD) of the respective curves. (B) Sulfide concentrations and OD600nm of sterile LB NaSH (1 mM) medium over time. The mean of three independent replicates is plotted. Error bars indicate the SD. (C) Schematic overview of sulfide oxidation and subsequent product formation by the reaction with oxygen at near-neutral pH. (D) Formation of sulfide oxidation products in sterile LB NaSH (1 mM) medium over the course of 90 min. Depicted are the mean values of three independent replicates. Error bars indicate the SD. Polysulfide formation measured as ‘area under the curve’ (AUC). (E) Growth curves of strains RN4220, RN4220 Δ*cst1*, and RN4220 Δ*cst1:cst1* in pre-incubated LB NaSH (1 mM) medium. Plotted are the mean values of three independent, biological replicates. Light-colored areas indicate the SD of the respective curves. (F) Growth curves of strain Newman, Newman Δ*cst1*, and Newman Δ*cst1:cst1* in pre-incubated LB NaSH (1 mM) medium. Data shows the mean values of three independent, biological replicates. Light-colored areas indicate the SD of each curve.

In the first 30 minutes, a rapid decrease in sulfide concentration and a simultaneous increase in polysulfide concentration was observed. After 60 minutes, sulfide was undetectable. The polysulfide concentration decreased moderately as elemental sulfur was formed, and between 60 and 90 min the sulfide, polysulfide, and sulfur concentrations were stable, suggesting an equilibrium in the sulfur redox chemistry. Additionally, thiosulfate and sulfite were formed only in negligible amounts. Given the rapid depletion of sulfide in the medium, we suspected that polysulfides, rather than sulfide, were responsible for the observed growth impairment. To confirm this, we repeated the growth experiment but pre-incubated the LB NaSH medium for 2 h before inoculation to ensure sulfide depletion and polysulfide generation. We also included the MSSA strain Newman and its respective Δ*cst1* mutant (Newman Δ*cst1*) in the experiment to exclude strain-specific variations. Like in the previous experiment, produced polysulfides (from 1 mM NaSH) prolonged the lag phase regardless of the strain background. Again, the prolongation of the lag phase was substantially more pronounced in RN4220 Δ*cst1* and Newman Δ*cst1*, compared to their respective wild types (Fig. 3E and F).

Due to the limited prevalence of the *sqr* gene in the *cst1* clusters of most staphylococci (and its absence in *cst2* clusters), a plasmid (pCQ11_cst1) was constructed that contains the *cst1* cluster of *S. aureus* Newman, purposefully excluding the *sqr* gene. This plasmid was introduced into RN4220 Δ*cst1* and Newman Δ*cst1* to evaluate the *cst*-cluster’s functioning without the *sqr* gene. The plasmid rescued the wild-type behavior of both strains (RN4220 Δ*cst1:cst1*, Newman Δ*cst1:cst1*) in the presence of generated polysulfides (Fig. 3E and F), thus not only confirming the importance of the *cst* cluster for growth under these conditions but also proving the *sqr* gene as non-essential for polysulfide detoxification.

Controls with other inorganic sulfur compounds (thiosulfate, sulfite) showed no effect on growth, confirming that polysulfides were the primary sulfur species affecting *S. aureus* growth (Fig. S4). In summary, polysulfides are sufficient to cause the observed growth inhibition in *S. aureus*, and *cst1* functions as a polysulfide detoxification cluster enabling faster re-initiation of bacterial growth.

### Effect of polysulfides on the growth behavior of *Staphylococcus* species

To examine wether the *cst*-encoded polysulfide detoxification is relevant to all staphylococci, we measured the growth effect of generated polysulfides on multiple strains of representative *Staphylococcus* species across the phylogeny (Fig. 4, Fig. S5). To determine a relative polysulfide tolerance level, we used the lag phase prolongation compared to the growth behavior of the negative control (Fig. 4A). 100% polysulfide tolerance corresponded to no difference in growth and 0% tolerance indicated that we observed no growth within 16 hours of incubation in medium containing generated polysulfides. Corroborating our previous results with *S. aureus*, we found that strains with *cst1* were significantly more tolerant to polysulfides than strains without the cluster, despite some variation between and within species (Fig. 4B and C). Strains with *cst1* showed moderate to high polysulfide tolerances of 35.26% to 88.31%, regardless of the *sqr* gene. Strains lacking the *cst1* gene cluster generally showed little to no polysulfide tolerance. This was most noticeable in *S. epidermidis* and *S. haemolyticus*, where we used wild-type strains naturally harboring or lacking the cluster. Overall, our analysis of various staphylococci clearly showed that the presence of *cst1* corresponds to higher levels of polysulfide tolerance. We concluded that polysulfide-containing environments have a substantial impact on staphylococcal growth and that *cst1* is crucial for polysulfide detoxification to less hazardous sulfite in staphylococci.

**Fig 4.**
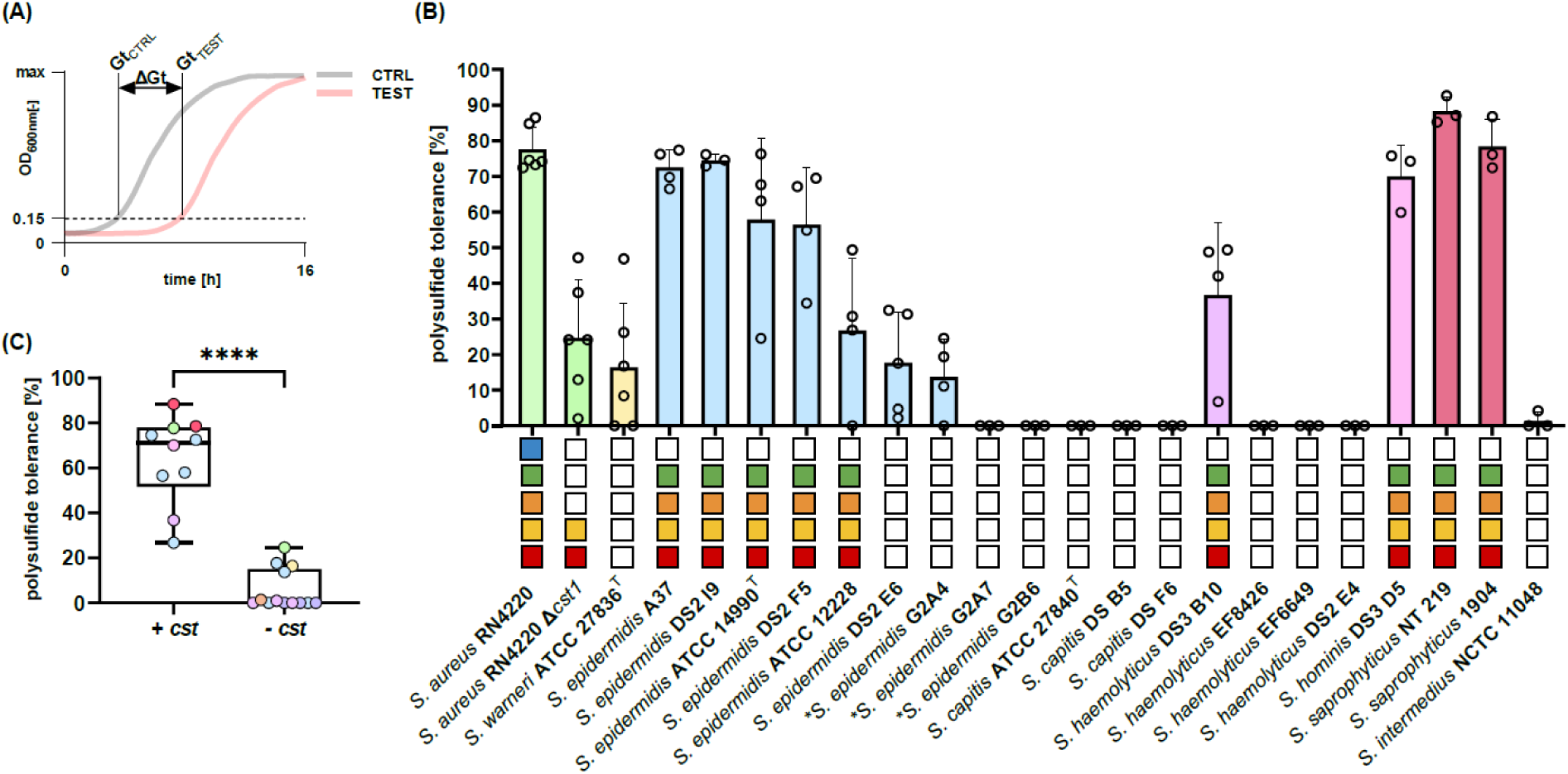
Polysulfide tolerance of different staphylococci. (A) Schematic illustration depicting the process of determining the relative polysulfide tolerance. The growth threshold (Gt) marks the time point where a bacterial culture reached OD600nm = 0.15. Depicted are the Gt value of the growth control (GtCTRL), the Gt of the test group (GtTEST), and the difference between GtCTRL and GtTEST (ΔGt). (B) Relative polysulfide tolerance of different staphylococci in LB containing generated polysulfides from 1 mM NaSH. Plotted are the mean values of at least three independent, biological replicates. Points show the data of the individual replicates. Squares denote the presence/absence of *cst* genes following the color scheme from Fig 1. (C) Comparison of the polysulfide tolerance of the analyzed staphylococci from (B) sorted by the presence (+ *cst*) or absence (- *cst*) of *cst1*. Depicted are box plots without outliers showing the median and the first and third quartile. Whiskers mark the minimum and maximum of the data range. Asterisks indicate statistical significance (\*\*\*\**P* ≤ 0.0001) from two-tailed Student’s t-tests. Plots in (B) and (C) follow the general color scheme from Fig 1 for different *Staphylococcus* species.

### Polysulfide tolerance of *S. aureus* strains with different *cst* cluster configurations

Having determined the polysulfide detoxifying function of the *cst1* cluster products, we investigated how SCC*mec*-carrying staphylococci benefit from an additional *cst2* cluster. First, we complemented the MSSA mutant strains RN4220 Δ*cst1* and Newman Δ*cst1*, with a plasmid (pCQ11_cst2) carrying the *cst2* from the MRSA strain COL. This rescued the wild-type behavior in both strains, verifying that *cst2* is fully functioning as a polysulfide detoxification cluster (Fig. S6). Notably, the Δ*cst1*:*cst2* strains behaved similarly to the Δ*cst1*:*cst1* strains, illustrating that the gene clusters are functionally synonymous. Thus, we suspected that the additional *cst2* might further improve polysulfide tolerance in MRSA compared to strains carrying only one *cst* cluster. Therefore, we tested the polysulfide tolerance of various MRSA and MSSA strains with different *cst* cluster configurations (Fig. 5, Fig. S7A). In general, we observed high polysulfide tolerance in *S. aureus* at low polysulfide concentrations (from 1 mM NaSH), independent of the strain background (MRSA or MSSA). Two MRSA strains naturally lacked *cst* clusters and showed no polysulfide tolerance, corroborating our previous results. With polysulfides generated from 3 mM NaSH, however, the polysulfide tolerance of the MSSA strains decreased significantly (Fig. 5A and B). Remarkably, MRSA strains still had high tolerance levels under these conditions, resulting in a significant difference between strains carrying one or two *cst* clusters (Fig. 5A and B).

**Fig 5.**
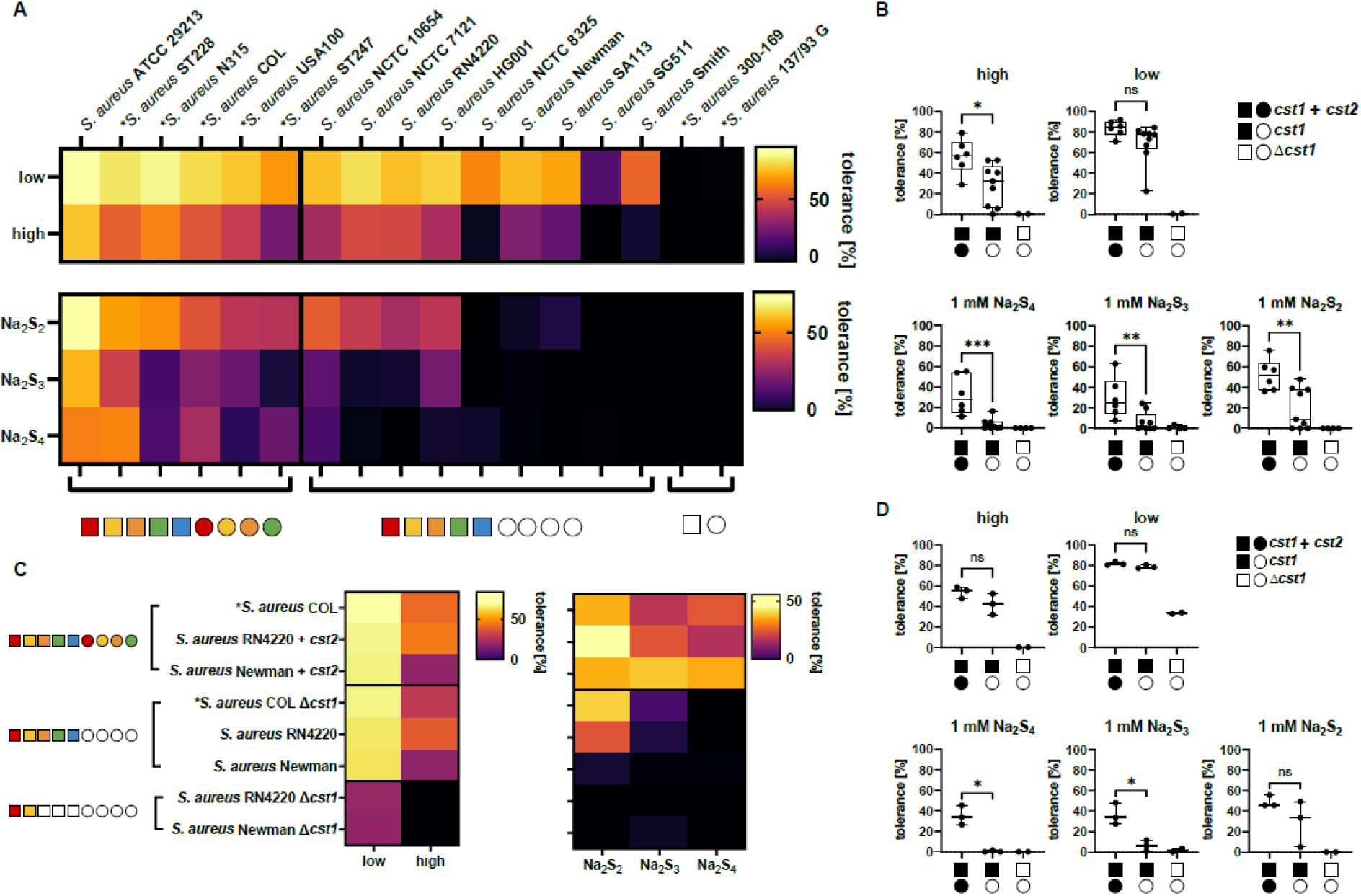
Impact of polysulfides on different *S. aureus* strains. (A) Heat maps displaying the relative polysulfide tolerance of multiple *S. aureus* wild-type strains at different polysulfide concentrations (“low”, generated from 1 mM NaSH; “high”, generated from 3 mM NaSH) and chain lengths (1 mM of Na2S2, Na2S3, or Na2S4) after 16 h of incubation. Depicted are the mean values of at least three independent, biological replicates. Squares and circles indicate the presence/absence of *cst1* or *cst2* genes, following the color scheme from Fig 1. MRSA strains are marked with asterisks in front of their strain designation. (B) Relative polysulfide tolerance of the tested strains from (A) grouped by harboring *cst1* + *cst2* (n=6), only *cst1* (n=9), or none (n=2). The box plots show the median and the first and third quartiles. Whiskers mark the minimum and maximum of the data range. Asterisks indicate statistical significance (ns = not significant; \**P* = 0.05 to 0.01; \*\**P* = 0.01 to 0.001; \*\*\**P* = 0.001 to 0.0001; \*\*\*\**P* ≤ 0.0001) from two-tailed Student’s t-tests. (C) Heat maps displaying the relative polysulfide tolerance of *S. aureus cst*-mutant strains and their respective wild types at different polysulfide concentrations (“low”, generated from 1 mM NaSH; “high”, generated from 3 mM NaSH) and chain lengths (1 mM of Na2S2, Na2S3, or Na2S4) after 16 h of incubation. Plotted are the mean values of at least three independent, biological replicates. Squares and circles indicate the presence/absence of *cst1* or *cst2* genes, following the color scheme from Fig 1. MRSA strains are marked with asterisks in front of their strain designation. (D) Relative polysulfide tolerance of the tested strains from (C) grouped by their cst configuration (*cst1* + *cst2* n=3, *cst1* n=3, and *Δcst1* n=2) and depicted as box plots. Shown are the median and first and third quartiles. Whiskers mark the minimum and maximum of the data range. Statistical significance from unpaired two-tailed Student’s t-tests is denoted as asterisks (ns = not significant; \**P* = 0.05 to 0.01; \*\**P* = 0.01 to 0.001; \*\*\**P* = 0.001 to 0.0001).

We hypothesized that the *cst2* cluster may improve MRSA fitness at higher polysulfide concentrations or in the presence of polysulfides with longer chain lengths. To this end, we repeated the previous experiment using polysulfides (1 mM) with defined chain lengths (Na2S2, Na2S3, and Na2S4). Now, the difference between MSSA and MRSA became even clearer (Fig. 5A, Fig. S7B). Again, strains lacking the *cst* cluster could not grow. The MSSA strains showed a heterogeneous behavior in the Na2S2 setup, with half the strains exhibiting medium tolerance, while the others failed to thrive. With increasing chain lengths MSSA tolerance steadily declined. In the Na2S4 setup, MSSA polysulfide tolerance was drastically limited, with only one of eight strains showing more than 10% tolerance. In contrast, MRSA strains had significantly higher tolerance levels to all three polysulfide chain lengths tested. We still observed that increasing the chain length affected MRSA tolerance to a varying degree. However, the mean tolerance of MRSA strains consistently exceeded that of the most tolerant MSSA strain. Even in the presence of 1 mM Na2S4, all MRSA strains grew. None showed less than 10% tolerance and two of six strains still exhibited over 50% tolerance.

To validate these results, we replicated the experiment with genetically modified strains, introducing a plasmid-located *cst2* (pCQ11_cst2) from COL into MSSA wild-type strains (RN4220 and Newman), and deleting *cst1* from MRSA COL (Fig. 5C). While the polysulfides generated from 1 mM or 3 mM NaSH induced only small, non-significant differences between the strains harboring one or two *cst* clusters (Fig. 5B), significant differences were observed when 1 mM Na2S3 and 1 mM Na2S4 were applied (Fig. 5D). Strains having both clusters (*cst1* and *cst2*) were significantly more tolerant than strains with only one cluster at polysulfide chain lengths of 3 or 4, regardless of the strain background and *cst-*cluster-configuration. Consequently, MSSA mutants with added *cst2* behaved similarly to MRSA wild-type strains (see also Fig. 5A for comparison), while the MRSA mutant COL carrying only *cst2* was phenotypically comparable to the MSSA wild-type strains. These results confirmed that the additional SCC*mec*-located *cst2* is largely responsible for the phenotypic polysulfide tolerance in MRSA. Moreover, the use of polysulfide standards demonstrated that the inhibitory effect is not entirely dependent on the polysulfide concentration but rather on the amount of sulfur bound in the reactive polysulfide form.

### Intraspecies competition between MSSA and MRSA under polysulfide stress

Based on the reported results, we hypothesized that the increased polysulfide tolerance of MRSA may enhance its competitive fitness against MSSA in polysulfide-containing environments. Therefore, we co-incubated MSSA strain RN4220 with either of two MRSA strains (COL and N315, representing SCC*mec* type I with Group B *cst2*, and SCC*mec* type II with Group A *cst2*, respectively) in a 99:1 OD600 ratio. We monitored their relative abundance over three days, with daily passaging into fresh medium containing 1 mM Na2S3. Without polysulfides, the MSSA strain remained at an overwhelming majority of ≥ 99% relative abundance throughout the experiment (Fig. 6), indicating that the MRSA failed to outcompete the other strain and invade the occupied environment. However, under polysulfide stress conditions, the MRSA strains were able to compete with the MSSA strain RN4220, establishing a substantial subpopulation of over 15% within the first 24 h of co-incubation (Fig. 6). After 48 hours, the MRSA strains dominated the co-cultures. After 72 hours, both MRSA strains represented over 98% of the co-cultures, completely reversing the initial ratio within three days. As expected, the MSSA strain Newman could not benefit from the polysulfide stress, and the strain was thus unable to outcompete the occupying RN4220 during the control experiment. This demonstrated that the SCC*mec*-located *cst2* enables MRSA to drastically outcompete MSSA in polysulfide-rich environments.

**Fig 6.**
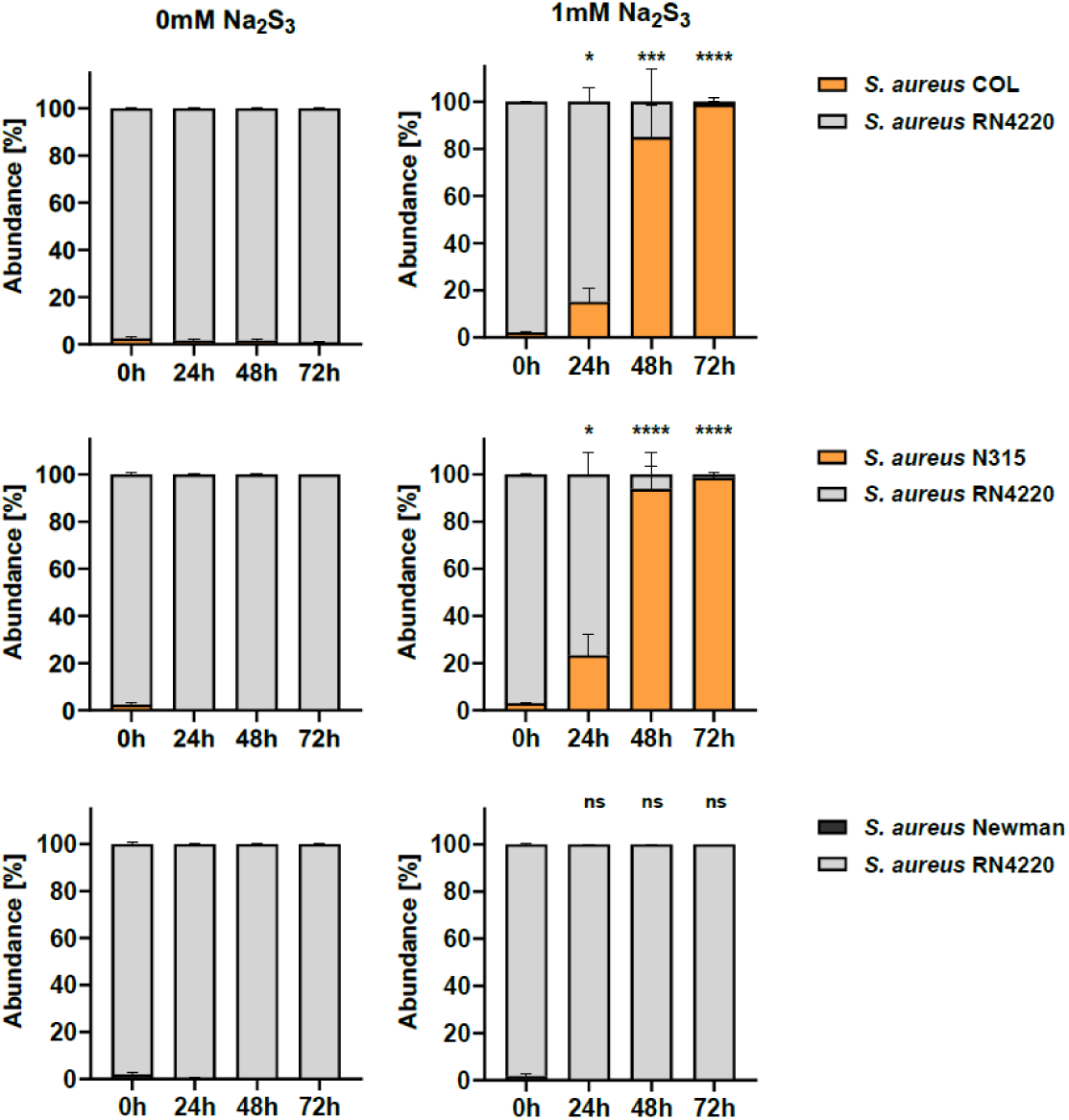
Competitive growth behavior of MRSA and MSSA (Newman) against the MSSA strain RN4220. Shown are relative strain abundances during serial passaging over the course of 72 h in medium with or without 1 mM Na2S3. Depicted are the mean values of three independent, biological replicates. Error bars indicate SD. Statistical significance from unpaired two-tailed Student’s t-tests is denoted as asterisks (ns = not significant; \**P* = 0.05 to 0.01; \*\**P* = 0.01 to 0.001; \*\*\**P* = 0.001 to 0.0001; \*\*\*\**P* ≤ 0.0001).

## Discussion

In this work, we have elucidated the ecophysiological role of the *cst* gene clusters in mediating polysulfide detoxification in staphylococci. Notably, we have shown that MRSA strains frequently harbor an additional, SCC*mec*-located *cst* cluster, namely *cst2,* which substantially increases polysulfide tolerance and provides deciding competitive fitness in polysulfide-rich environments.

The widespread, heterogeneous distribution of the *cst1* gene cluster in staphylococci emphasizes its importance as an adaptation to specific, sulfide-containing habitats. Although *Staphylococcus* species are common colonizers of the human body, not all host-associated staphylococci will encounter (poly-)sulfides, as the conditions necessary for generating these compounds differ drastically among different body sites. While strains inhabiting mucosal surfaces presumably face these stressors regularly, other strains colonizing, e.g., the skin are much less exposed to (poly-)sulfides. Thus, possessing *cst* gene clusters may serve as an adaptation to mucosal habitats. Moreover, since various forms of inflammation are linked to increased sulfide levels (31), the *cst* clusters may be crucial for pathogenic or infection-associated strains.

In contrast to the overall abundance of the *cst1* cluster, the *sqr* gene was predominantly restricted to *S. aureus* and was never present in *cst2* clusters. This suggests a particular ecological function of the SQR beyond the general role of *cst1*. In support of this, we observed that *cst* clusters provide fully functional polysulfide detoxification without *sqr*, demonstrating that TauE, CstA, and CstB are sufficient to confer polysulfide tolerance to staphylococci. Thus, we propose to include polysulfide detoxification without SQR as an extension to the current model of the gene cluster’s function. Without the need for the initial sulfide oxidation by SQR, polysulfides may directly interact with CstA, which transfers individual thiol groups onto a suitable cellular sulfur acceptor, i.e. a low-molecular-weight thiol (LMW-SH). The resulting LMW-SSH may be handled by CstB, which oxidizes the persulfide sulfur with the consumption of O2 and releases sulfite, which is finally transported out of the cell by TauE (Fig. S8). Longer polysulfide chains may increase stress due to the increased amount of reactive thiol groups that need to be transferred to CstA, which is supported by our results with the polysulfide standards. Regarding the ecological function of SQR in *S. aureus*, it remains elusive how its well-characterized role in the sulfide-to-thiosulfate conversion with concomitant quinone reduction (18) translates into a trait that is beneficial for this species.

While great advances have been made to elucidate various effects of H2S on bacterial physiology, little is known about the impact of other reactive sulfur compounds (32). Our results demonstrate that polysulfides are highly toxic to staphylococci, matching or exceeding H2S toxicity. Detoxification by *cst1*-encoded enzymes reduces exposure time to harmful polysulfide concentrations, contributing to the observed polysulfide tolerance. The SCC*mec*-located *cst2* cluster provides additional detoxification capacity, enhancing the tolerance to high polysulfide stress. Our bioinformatics analysis demonstrated that *cst2* is widespread amongst MRSA strains, spanning key SCC*mec* types and multiple epidemiologically relevant strains (27, 28). To the best of our knowledge, this constitutes *cst2* as the most abundant SCC*mec*-located gene cluster (aside from the SCC*mec*-defining *mec* and *ccr* complexes). This strongly suggests that MRSA benefits from *cst2*-conferred tolerance to proliferate in (poly-)sulfide-rich niches, like the nose and, in particular, the gut (33, 34, 4). Moreover, this may be especially important during events of dysbiosis and inflammation, where levels of reactive sulfur species are elevated (5). In turn, this may be relevant in clinical settings, where prolonged hospitalizations and medical predispositions often cause steady levels of dysbiosis and inflammation (e.g., long-term care facilities, dialysis, post-surgery, and ICU admission) (35, 36). In support, we found that the *cst2* cluster is prominent in SCC*mec* types strongly associated with HA-MRSA (27, 28).

Since our co-incubation experiments demonstrated that the *cst2* cluster effectively provides MRSA with a substantial advantage in intraspecies competition against already established MSSA populations during polysulfide stress, the additional gene cluster might act as a colonization factor enabling MRSA to invade the human microbiome. This might be especially relevant because the microbiome of mucosal surfaces itself is a constant source of considerable (poly-)sulfide concentrations (6, 37, 7, 8, 38). Polysulfide tolerance may therefore select for MRSA over less tolerant staphylococci, particularly during states of disease.

Overall, our work shows that polysulfide stress is a biologically relevant factor for staphylococci that may contribute to the spread of *cst2*-containing SCC*mec* and MRSA. Thus, we suggest that the SCC*mec*-located *cst2* cluster is a critical factor facilitating MRSA proliferation within the human microbiome.

## Methods

### Bioinformatics analyses

The *cst* gene cluster was identified using a modified version of the HMSS2 tool (22) by extending the library to include the proteins CstR, CstA and MecA. The Distribution of the *cst* gene cluster and *mecA* were visualized in a maximum likelihood phylogeny. For detailed information, see supplementary material and methods.

The presence of *cst2* in SCC*mec* cassettes was determined using BLAST for a list of representative MRSA strains (Table S2), compiled from sources listing reference strains for SCC*mec* typing (24–26) and epidemiologically relevant strains (27, 41, 28). To generate the phylogram of staphylococcal *cst* gene clusters, all thereby found SCC*mec*-located *cst2* clusters and representative genomic staphylococcal *cst1* gene clusters detected with the HMSS2 tool were aligned using EMBL-EBI Clustal Omega (42). The phylogenetic tree was calculated using IQTree 1.6.12 (43) and visualized with FigTree v1.4.4. Representative SCC*mec* cassettes carrying *cst2* were aligned using EMBL-EBI Clustal Omega (42) and visualized using clinker (44).

To determine *cst2* distribution in publicly available complete MRSA genomes, *cst2* from N315 or COL were used as query sequences for *cst2* Group A and B, respectively, in a BLAST genome search of the NCBI *S. aureus* (taxid:1280) complete genome database. Results were filtered for >90% identity to exclude *S. aureus cst1* and the respective other *cst2* group. Shortened Group B *cst2* (*cstB2*) was identified by query cover. The total number of MRSA genomes was determined by a similar BLAST genome search using *S. aureus* N315 *mecA* as query sequence and all strains featuring a full-length *cst2* were filtered for the presence of *mecA* by list comparison with the results of this search.

### Strains and plasmids

All primers, plasmids, and strains used in this study are listed in Table S7-9. The generation of *S. aureus* Δ*cst1* mutants and the introduction of plasmid-located *cst1* and *cst2* into *S. aureus,* as well as the detection of the *cst* gene cluster in *Staphylococcus* spp. are described in supplementary material and methods.

### Standard growth conditions

All strains were cultivated in Lysogeny Broth (LB, 10 g/l NaCl) and incubated at 37 °C and 120 rpm agitation, if not stated otherwise.

### Generation of polysulfides in LB medium

To generate different concentrations of polysulfides (low, high), LB medium was supplemented with different concentrations of NaSH (1 mM and 3 mM, respectively) and incubated for 2 h at 37 °C with 120 rpm agitation to ensure sufficient polysulfide generation.

### Growth experiments with sulfide and polysulfides

To study the impact of sulfur species on the growth of different staphylococci, 99 µl of generated polysulfides (see above), 1 mM NaSH, 1 mM Na2Sn standards (Na2S2, Na2S3 or Na2S4), 1 mM S2O3^2−^, 1 mM SO3^2−^, or 1 mM SO4^2−^ dissolved in LB medium were transferred into a 96-well plate, inoculated with 1%(v/v) of an overnight grown culture (OD600nm adjusted to 0.3 prior to inoculation), and sealed with gas-permeable MICRONAUT sealing foil (sifin diagnostic). The optical density was measured every 5 min for 16 h at 37 °C using a Tecan infinite 200 PRO (Tecan).

The lag phase prolongation was used to determine a relative polysulfide tolerance level compared to the growth behavior of the negative control. 100% polysulfide tolerance corresponded to no difference in growth and 0% tolerance denoted that we did not observe growth within 16 h of incubation in medium containing generated polysulfides. To determine the time point where cultures re-initiated growth, a growth threshold (Gt) value at which cultures reached OD600nm = 0.15 (marking the beginning of the log phase) was defined (see also Fig. 4A). The difference (ΔGt) between the Gt value under polysulfide stress (GtTEST) and the Gt value of the negative control (GtCTRL) was then used to calculate the polysulfide tolerance using equation (1).

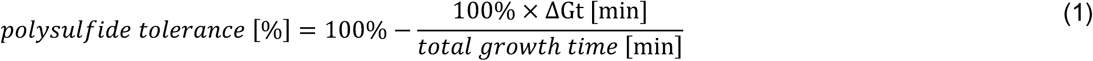

### Competition Assay

Overnight cultures of MSSA (RN4220, Newman) and MRSA (COL, N315) strains were adjusted to OD600 = 1.5 and mixed at an OD600 ratio of 99:1 (MSSA/MRSA, MSSA/MSSA) to a final volume of 1.5 ml. LB medium with or without 1 mM Na2S3 was inoculated with 1% (v/v) of the mixture and incubated for 24 h at 37 °C with 120 rpm agitation. After 24 h and 48 h, fresh medium with or without 1 mM Na2S3 was inoculated with 1%(v/v) of the competition mixture. For each time point (0 h, 24 h, 48 h, 72 h), the competition mixture was serially diluted in 0.9% NaCl solution and 10µl of it was spotted on LB plates for total cell numbers and LB plates with antibiotic for strain-specific selection (10 µg/ml erythromycin, 2.5 µg/ml oxacillin, or 10 µg/ml chloramphenicol, respectively, see Table S11). After 24 h of growth, the colony-forming units (CFUs) per ml were determined for each strain, and the MSSA/MRSA and MSSA/MSSA ratios were determined.

### Determination of sulfur species

To determine the concentrations of HS^-^ and its oxidation products in medium over time, LB medium was substituted with 1 mM NaSH and incubated for 2 h at 37 °C with 120 rpm agitation. At different time points, samples were taken to quantify the sulfide species. Thioles were quantified after derivatization with the fluorescent dye monobromobimane as previously described (45). S8 was colorimetrically determined after treatment with cyanide as described elsewhere (46). See supplementary material and methods for a detailed description.

### Statistical analysis

Statistical analysis of data was performed using GraphPad Prism 10.2.0. If not stated otherwise, statistical significance was determined in unpaired two-tailed Student’s *t*-tests with a 95% confidence interval against the respective standard or negative control. All experiments were repeated at least three times with biologically independent replicates. All graphs show the mean of at least three biological replicates with error bars denoting the SD. Statistical significance was denoted as not significant (ns); \**P*=0.05 to 0.01; \*\**P*=0.01 to 0.001; \*\*\**P*=0.001 to 0.0001; \*\*\*\**P*≤0.0001.

## Data availability

All raw data for graphs are supplied in source data files. Source data are provided with this paper.

## Code availability

HMSS2 program files are available at https://github.com/TSTanabe/HMSS2.

## Acknowledgements

We thank Jana Rohe and Kainat Qureshi for technical assistance, and Prof. Dr. Gabriele Bierbaum and her research group for supplying us with staphylococcal isolates from their remarkable collection. We would also like to thank Dr. Stefania De Benedetti for additional proofreading. This work was mostly funded by the Jürgen Manchot Foundation and the German Center for Infection Research (DZIF). Tomohisa Sebastian Tanabe acknowledges funding from the Austrian Science Fund (FWF) [doi.org/10.55776/COE7].

## Ethics declarations

The authors declare no competing interests.

## Notes

### Competing Interest Statement

The authors have declared no competing interest.

