## Supplemental Materials for "A widespread SCC*mec*-located gene cluster protects methicillin-resistant *Staphylococcus aureus* against toxic polysulfides"

Running head: *cst2* confers high polysulfide tolerance to MRSA

* Present address: Fabian Grein, biovis Diagnostik MVZ GmbH, Limburg-Eschhofen, Germany

**The Supplemental Materials include:**

- Supplementary Material and Methods
- Figure S1: Current model of the detoxification of $H_2S$ in *S. aureus*
- Figure S2: Plasmid Map of pATCC29213 from *S.aureus* strain ATCC29213
- Figure S3: Alignment of *S. aureus cst1* and SCC*mec* cassettes featuring a *cst2* cluster
- Figure S4: Growth behavior of *S. aureus* wild type and Δ*cst1* mutants in LB medium supplemented with 1 mM $S_2O_3^{2-}$ /$SO_3^{2-}$
- Figure S5: Growth curves of different staphylococci in LB medium containing generated polysulfides
- Figure S6: Growth behavior of *S. aureus* wild type and mutants in LB medium containing generated polysulfides
- Figure S7: Growth curves of different *S. aureus* strains in LB medium containing generated polysulfides/ 1 mM polysulfide standards
- Figure S8*:* Postulated model of SQR-independent polysulfide detoxification in *S. aureus*
- Table S1: GenBank accession numbers for the construction of the phylogenetic tree and the distribution of *cst* gene clusters and *mecA* in this study
- Table S2: list of MRSA reference strains and epidemiologically relevant strains for SCC*mec* typing
- Table S3: List of *S. aureus* with *mecA*-containing genomes by screening the NCBI complete genome database
- Table S4: List of *S. aureus* genomes containing group A *cst2*
- Table S5: List of *S. aureus* genomes containing full-lenght group B *cst2*
- Table S6: List of *S. aureus* genomes containing shortened version of Group B *cst2*
- Table S7: Oligonucleotides used in this study
- Table S8: Plasmid used in this study
- Table S9: Strains used in this study
- Table S10: CstR reference proteins to build the CstR hidden Markov model in this study
- Table S11: CstA reference proteins to build the CstA hidden Markov model in this study
- Table S12: MecA reference proteins to build the MecA hidden Markov model in this study
- References

#### Supplementary Material and Methods

##### Bioinformatics analyses

The HMSS2 (1) tool identified sulfur metabolism-related genes and analyzed operon structures in genome and metagenome assemblies based on hidden Markov models (HMM). The HMSS2 library already contained HMM for the *cst* gene products proteins TauE, CstB known as SDOIII-cstA, and SQRII. The library was extended to incorporate the missing proteins CstR and CstA from the gene cluster by constructing an HMM based on aligned staphylococcal CstR and CstA protein sequences obtained from UniProt database or NCBI protein database (Table S10 and S11). To detect the presence of SCC*mec*, an additional HMM was generated for the protein MecA (Table S12). 230 genomic assemblies of members of the family Staphylococcaceae were downloaded from NCBI RefSeq or GenBank and then searched for *cst* genes and *mecA* using HMSS2 (Table S1). For phylogenetic clustering, nucleotide FASTA files were downloaded from NCBI. Open reading frames were determined with prodigal (2). RNA polymerase subunits were searched with the hidden Markov models TIGR02013 for *rpoB* and TIGR02386 for *rpoC* with the cutoffs from TIGRFAMs. The phylogenetic tree was inferred from concatenated alignments of RNA polymerase proteins RpoB and RpoC. Protein sequences were concatenated and subsequently aligned using MAFFT with default setting. The alignment was trimmed using BMGE (entropy threshold = 0.95, minimum length = 1, matrix = BLOSUM65). Alignments were then used for maximum likelihood phylogeny inference using IQ-TREE v1.6.12. The best-fitting model of sequence evolution was selected using ModelFinder. Branch support was calculated by SH-aLRT (2000 replicates), aBayes (2000 replicates), and ultrafast bootstrap (2000 replicates). The phylogenetic tree and the abundance of *cst* gene clusters and *mecA* were visualized with iTOL version 5 (3).

##### Generation of *S. aureus* ∆*cst1* mutants

*S. aureus* RN4220 ∆*cst1* was generated using a previously described technique with the plasmid pMAD (4). To generate pMAD_*cst1*, DNA fragments were PCR-amplified from *S. aureus* RN4220 genomic DNA using the primers ∆cstA1_SalI_for and ∆cstA1_EcoRI_rev, and ∆sqr_EcoRI_for and ∆sqr_NocI_rev, digested with *Eco*RI, ligated (T4 DNA ligase, New England Biolabs (NEB), Frankfurt am Main, Germany) and PCR-amplified using primers ∆cstA1_SalI_for and ∆sqr_NocI_rev. This fragment was cloned into pMAD using *Sal*I and *Noc*I and pMAD_*cst1* was transformed into *E. coli* TOP10

(Invitrogen, Thermo Scientific, Schwerte, Germany). The chloramphenicol resistance cassette (CAT; chloramphenicol acetyltransferase) was amplified from the plasmid pC194 using the primers CAT_for and CAT_rev and cloned into pMAD_*cst1* using *Eco*RI. The functional deletion of *cst1* in *S. aureus* RN4220 was performed as previously described via double homologous recombination (4) and verified with PCR using the primers Check_Δcst1_for and Check_Δcst1_rev.

*S. aureus* Newman Δ*cst1* and *S. aureus* COL Δ*cst1* were created using phage transduction (phage Φ11) by transferring the genomic region containing the deletion genotype Δ*cst1* from *S. aureus* RN4220 Δ*cst1* to the target strains *S. aureus* Newman and *S. aureus* COL as previously described (5).

**Introduction of plasmid-located *cst1* and *cst2* into *S. aureus***

For the construction of pCQ11_*cst1* and pCQ11_*cst2*, *cst1* was amplified from *S. aureus* Newman genomic DNA using the primers tauE1_NheI_for and cstB1_AscI_rev. *cst2* was amplified from *S. aureus* COL genomic DNA using the primers cstB2_NheI_for and tauE2_AscI_rev. The resulting fragments were cloned into pCQ11-mNeonGreen (6) using *Nhe*I and *Asc*I under simultaneous removal of *mNeongreen*. pCQ11_*cst1* and pCQ11_*cst2* were transformed into electrocompetent *S. aureus* RN4220, *S. aureus* RN4220 Δ*cst1*, *S. aureus* Newman and *S. aureus* Newman Δ*cst1* as previously described (7) using *Escherichia coli* 10β (NEB, Frankfurt am Main, Germany) or *E. coli* DC10β as shuttle strain.

**Detection of *cst* in *Staphylococcus* spp.**

Genomic DNA of *Staphylococcus* spp*.* was extracted using the NucleoSpin Microbial DNA kit (Macherey-Nagel, Düren, Germany) after pretreatment with 50 µg/ml lysostaphin. To detect the *cst* cluster, the DNA fragment was PCR-amplified, using the primers Staph_cstB_for1 and Staph_tauE_rev1 or Staph_tauE_for2 and Staph_cstB_rev2, and the presence of *cst* genes was determined by agarose gel electrophoresis.

**Determination of sulfur species**

Detection and quantification of thioles like $HS^-$, $^-SS_nS^-$, $S_2O_3^{2-}$ and $SO_3^{2-}$ was performed by using monobromobimane (8). Briefly, each sample was incubated with monobromobimane and the derivatized fluorescent sulfur species were subjected to HPLC analysis. HPLC was performed on an Agilent 1260 Infinity II System (Agilent Technologies) equipped with an FLD (Ex.: 380 nm; Em.: 480 nm) detector.

Separation was achieved on a LiChroCHART C8 (4 x 250 mm, 5 µm; Merck) with a flow rate of 0.5 ml/min at 30 °C. After short column acidification using 3 min of 0.25% acetic acid, sulfur species were eluted in a three-step linear gradient of MeOH (0 – 30% over 14 min, 31 – 55% over 23 min, 56 – 100% over 1 min). To determine the absolute concentrations of every sulfur species, monobromobimane fluorescence was quantified against respective standard curves using OpenLab CDS software version 2.6.

$S_8$ was colorimetrically determined by using cyanide (9). Briefly, each sample was centrifuged and the resulting pellet was washed and finally resuspended in 0.2 M sodium cyanide solution. After 10 min incubation at 100 °C and subsequent addition of 0.9 M ferric nitrate reagent in 22% $HNO_3$, the extinction at 460 nm was measured and quantified using a 0 - 3 mM NaSCN standard curve.

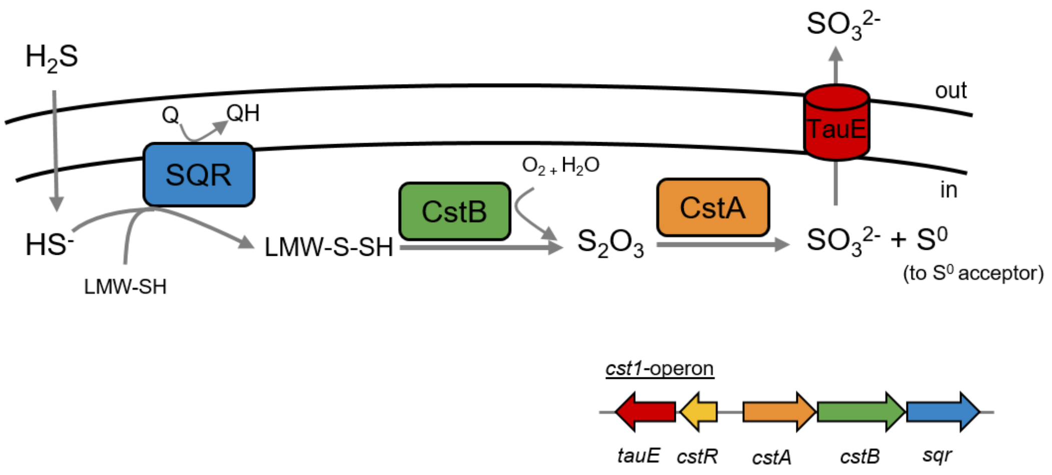


**Figure S1: Current model of the detoxification of $H_2S$ in *S. aureus*.** $H_2S$ crosses the cytoplasmic membrane and exists as $HS^-$ Anion. The $HS^-$ Anion is oxidized by the sulfide:quinone oxidoreductase (SQR, blue) in an initial two-electron oxidation step and the sulfane sulfur ($S^0$) is transferred to a low-molecular-weight thiol (LWM-SH), resulting in LMW persulfides (LMW-SSH). In the presence of oxygen ($O_2$), the sulfur dioxygenase (CstB, green) generates thiosulfate ($S_2O_3$) from the LMW-SSH. Subsequently, the sulfur transferase (CstA, orange) transfers the $S^0$ of $S_2O_3$ to a cellular sulfur acceptor, concurrently releasing sulfite ($SO_3^{2-}$), which is exported out of the cell by the sulfite transporter TauE (red) (Adapted from Shen *et al.* (2015) (10)).

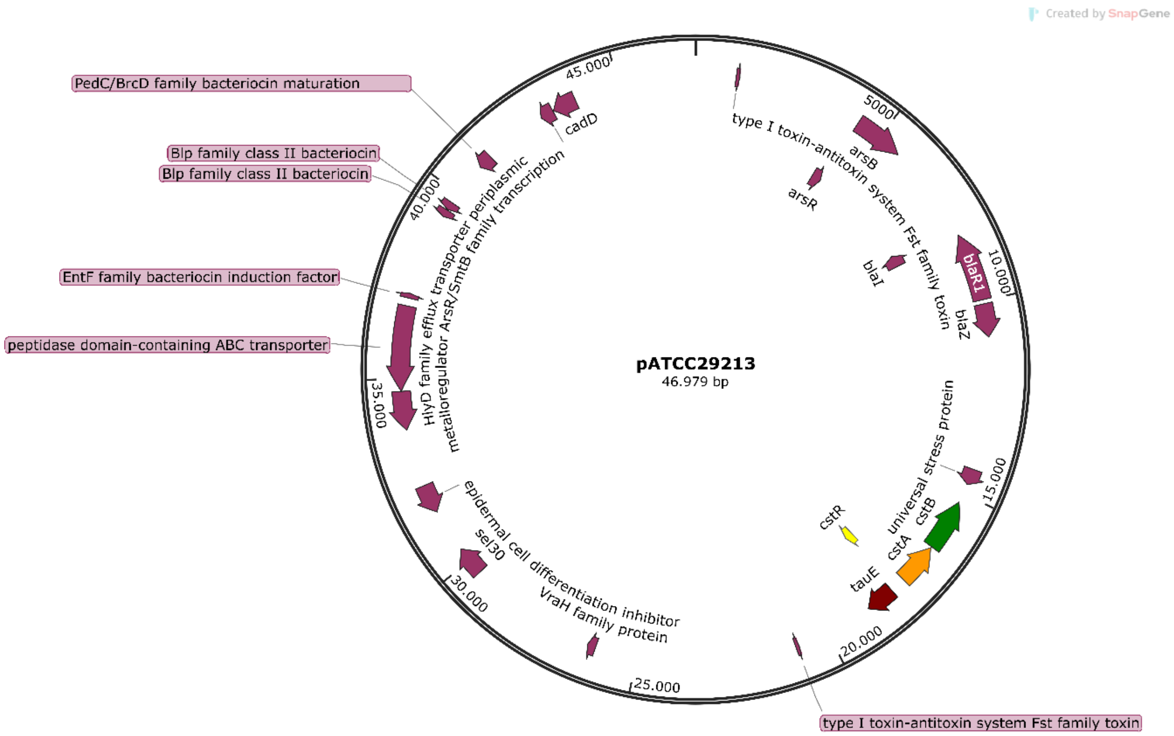


**Figure S2: Plasmid Map of pATCC29213 (NCBI GenBank: CP094858.1) from the *S. aureus* strain ATCC 29213 (NCBI GenBank: CP094857.1)**. Potential virulence factors and *cst2* genes are highlighted

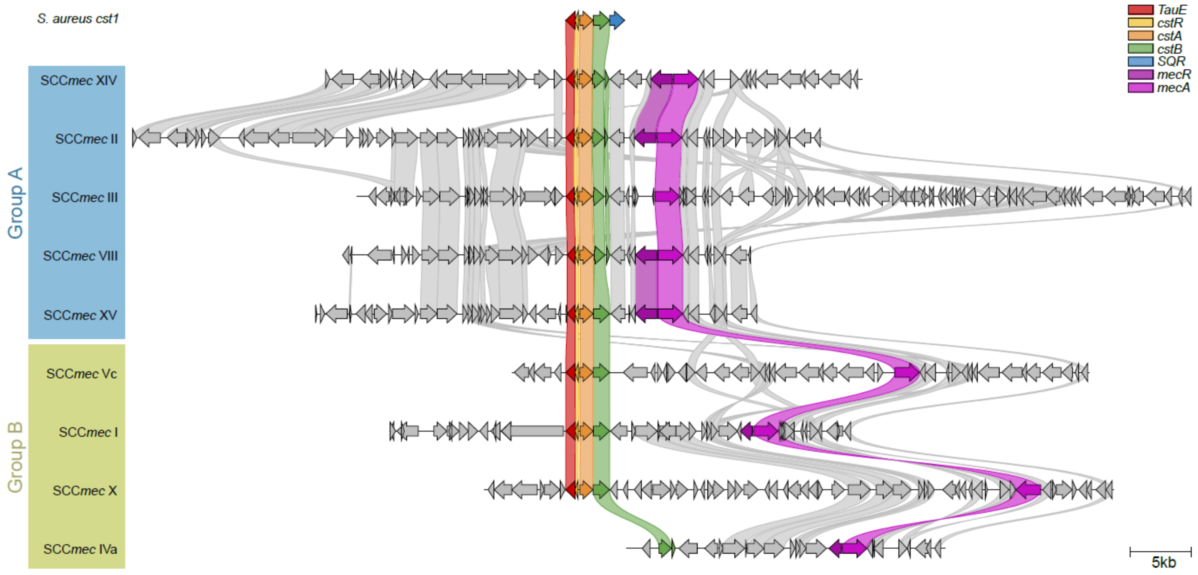


**Figure S3: Alignment of *S. aureus cst1* and SCC*mec* cassettes featuring a *cst2* cluster**. Connections denote annotated genes with ≥ 30% identity. *cst* genes, *mecA* and *mecR* are highlighted.

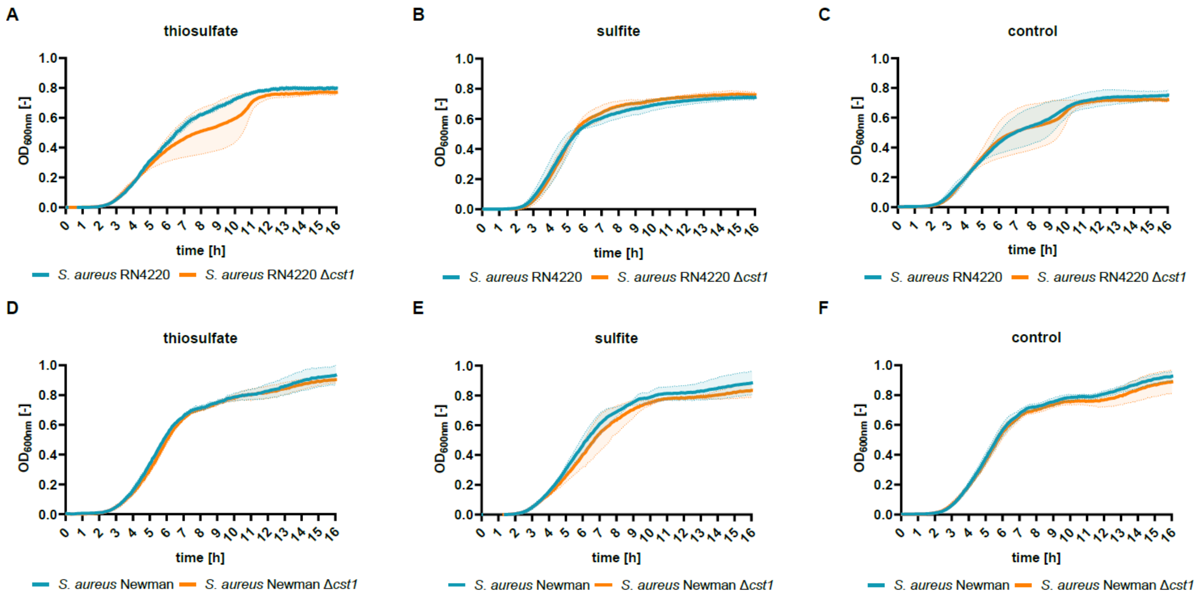


**Figure S4: Growth behavior of *S. aureus* wild type and Δ*cst1* mutants in LB medium supplemented with 1 mM $S_2O_3^{2-}$ /$SO_3^{2-}$.** Growth curves of the strains A) RN4220 and RN4220 Δ*cst1* or D) Newman and Newman Δ*cst1* in LB medium containing 1mM $S_2O_3^{2-}$ (thiosulfate). Growth curves of the strains B) RN4220 and RN4220 Δ*cst1* or E) Newman and Newman Δ*cst1* in LB medium containing 1mM $SO_3^{2-}$ (sulfite). In C) and F) the growth control of RN4220 and RN4220 Δ*cst1* or Newman and Newman Δ*cst1* in LB medium are depicted. The Data shows the mean values of at least three independent, biological replicates. Light-colored areas illustrate the SD of the respective curves.

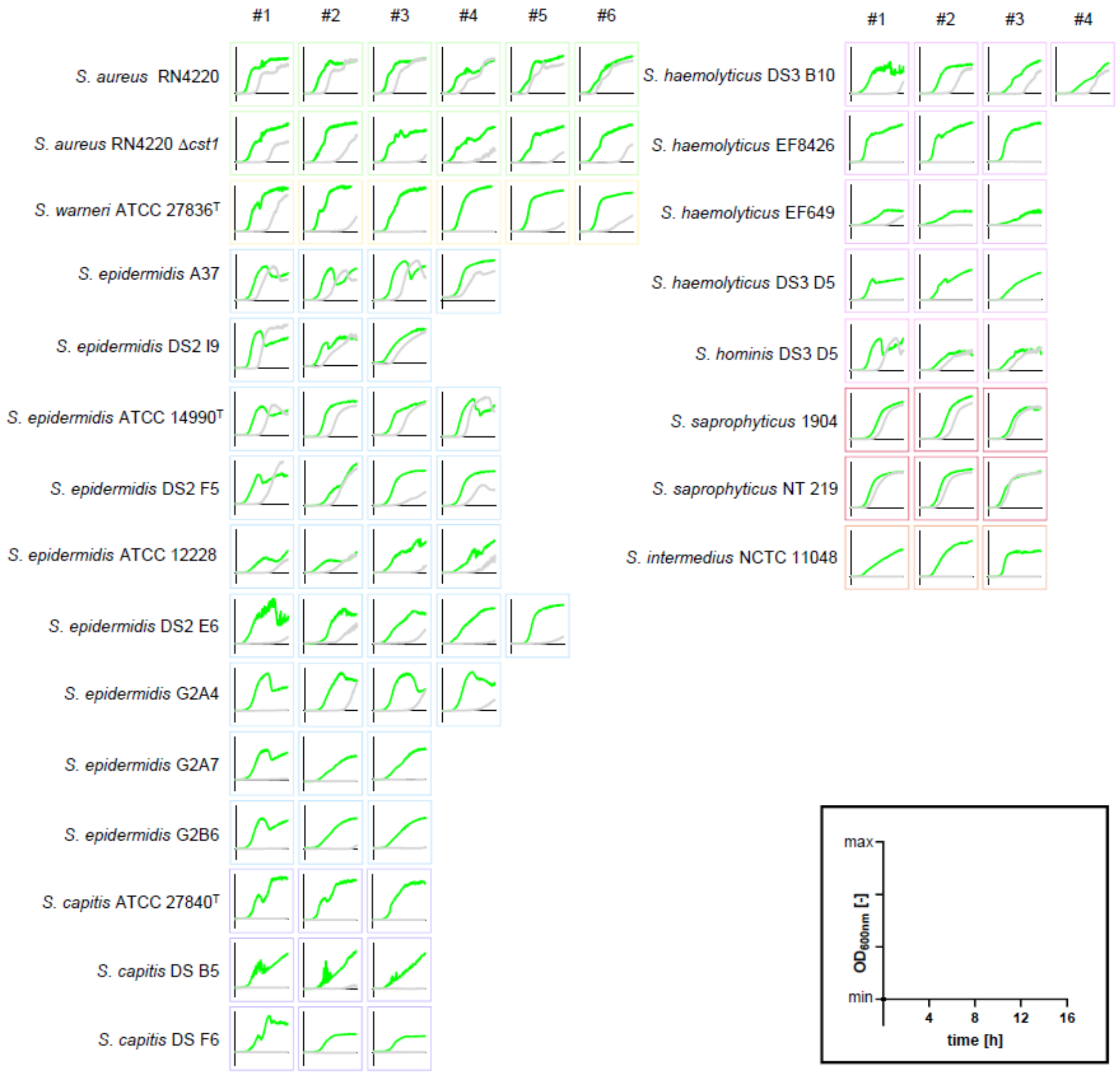


**Figure S5: Growth curves of different staphylococci in LB medium containing generated polysulfides**. Depicted are all biological replicates for each strain. Data contains the growth in polysulfide-containing LB medium (generated from 1 mM NaSH, light gray) and the control in LB medium without supplementation (green). The y-axis represents the $OD_{600nm}$ with strain-specific maximum, x-axis represents the incubation time with a maximum of 16h. The frames follow the general color code from Fig. 1 for different *Staphylococcus* species

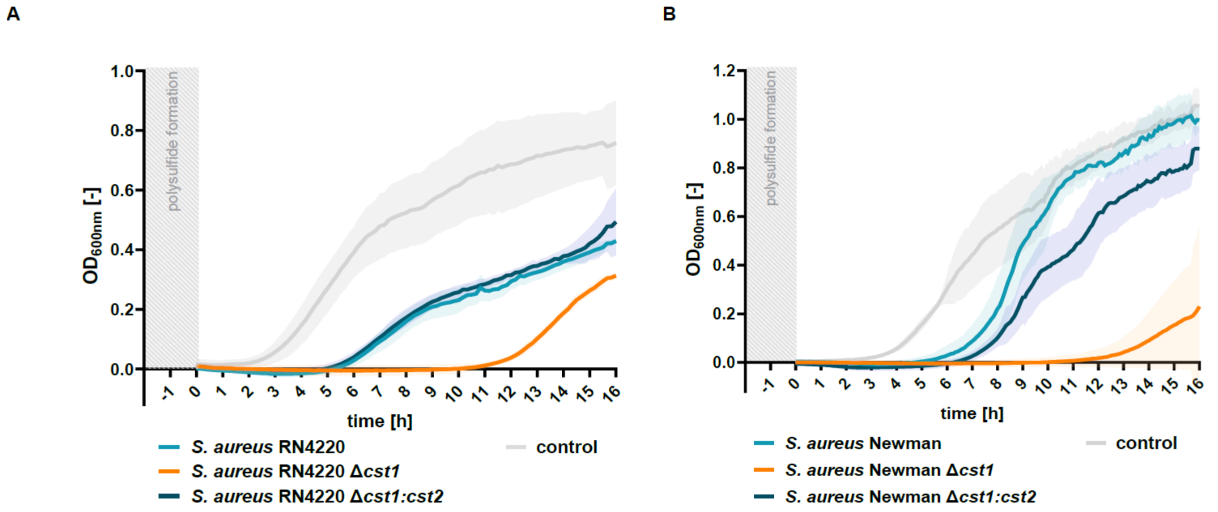


**Figure S6: Growth behavior of *S. aureus* wild type and mutants in LB medium containing generated polysulfides.** A) Growth curves of strain RN4220, RN4220 Δ*cst1*, and RN4220 Δ*cst1:cst2* in pre-incubated LB NaSH (1mM) medium. Plotted are mean values of three independent, biological replicates. Light-colored areas illustrate the SD of the respective curves. B) Growth curves of strain Newman, Newman Δ*cst1*, and Newman Δ*cst1:cst2* in pre-incubated LB NaSH (1mM) medium. Data shows the mean values of three independent, biological replicates. Light-colored areas illustrate the SD of the respective curves.

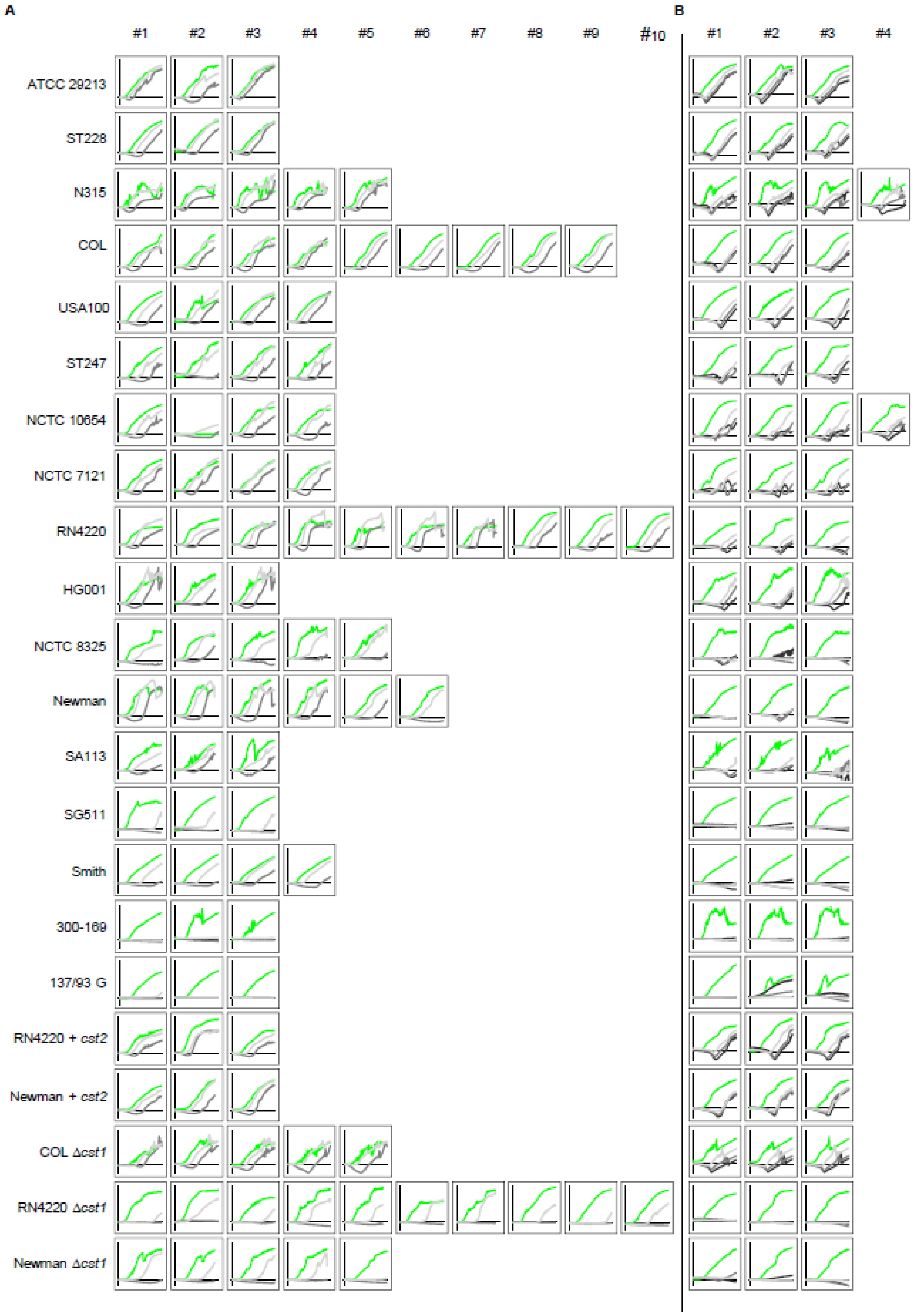


**Figure S7: Growth curves of different *S. aureus* strains in LB medium containing generated polysulfides/ 1 mM polysulfide standards.** Depicted are all biological replicates for each strain. A) Growth in LB medium containing generated polysulfides (2h pre-incubation of 1 mM NaSH (low, light gray) or 3 mM NaSH (high, dark gray) in LB medium). B) Growth in LB medium supplemented with 1 mM polysulfide standards ($Na_2S_2$ light gray, $Na_2S_3$ dark gray, $Na_2S_4$ black). Growth control in LB medium (green). Y-axis represents the $OD_{600nm}$ with strain-specific maximum, x-axis represents the incubation time with a maximum of 16h.

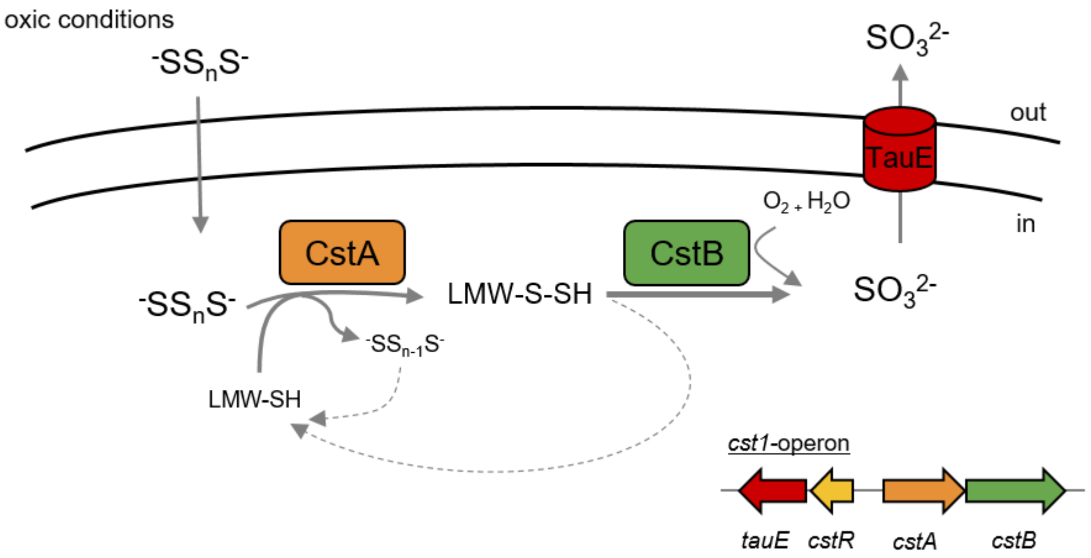


**Figure S8: Postulated model of SQR-independent polysulfide detoxification in *S. aureus.*** The polysulfide anion ($^-SS_nS^-$) crosses the cell membrane. In a first step the sulfur transferase (CstA, orange) transfers a sufane sulfur ($S^0$) from $^-SS_nS^-$ to an LMW thiol (LWM-SH), resulting in LMW persulfides (LMW-SSH). Subsequently, the sulfur dioxygenase (CstB, green) oxidizes the LMW-SSH with oxygen ($O_2$) and water ($H_2O$) to sulfite ($SO_3^{2-}$). In this process, the LMW thiol pool is regenerated and the polysulfide anion is reduced by one thiol group ($^-SS_{n-1}S^-$) and both are available for further oxidation. In the last step, $SO_3^{2-}$ is exported out of the cell by the sulfite transporter TauE (red).
